# Molecularly informed analysis of histopathology images using natural language

**DOI:** 10.1101/2025.07.14.664402

**Authors:** Moritz Schaefer, Kalin Nonchev, Animesh Awasthi, Jake Burton, Viktor H Koelzer, Gunnar Rätsch, Christoph Bock

## Abstract

Histopathology refers to the microscopic examination of diseased tissues and routinely guides treatment decisions for cancer and other diseases. Currently, this analysis focuses on morphological features but rarely considers gene expression information, which can add an important molecular dimension. Here, we introduce SpotWhisperer, an AI method that links histopathological images to spatial gene expression profiles and their text annotations, enabling molecularly grounded histopathology analysis through natural language. Our method outperforms pathology vision-language models on a newly curated benchmark dataset, dedicated to spatially resolved H&E annotation. Integrated into a web interface, SpotWhisperer enables interactive exploration of cell types and disease mechanisms using free-text queries with access to inferred spatial gene expression profiles. In summary, SpotWhisperer analyzes cost-effective pathology images with spatial gene expression and natural-language AI, demonstrating a path for routine integration of microscopic molecular information into histopathology.

## 1. Introduction

Histopathology is fundamental for clinical diagnostics and guides treatment decisions across a wide range of diseases. For example, most cancers are diagnosed through manual examination of macroscale morphological features in hematoxylin and eosin (H&E) stained tumor specimens under a microscope. Despite the low cost of this assay, the analysis is labor-intensive and requires trained pathologists.

A key goal of computational pathology is to make histopathology image analysis more accessible. Vision-language models (VLMs), which interpret images through natural-language queries, allow training and inference on H&E images annotated with broad disease phenotypes and macroscopic features (e.g., “tumor”, “inflamed tissue”). As a result, these models tend to focus on macroscale morphology and are at risk of overlooking the molecular and microanatomical features needed to capture more fine-grained aspects of tissue biology and pathophysiology.

By contrast, modern biomedical research increasingly focuses on molecular readouts such as gene expression profiling, which can provide mechanistic insight and enable fine-grained characterization of biological states. Spatial transcriptomics offers a way to obtain molecular information at high spatial resolution across thousands of genes simultaneously. However, the high cost and substantial technical and bioinformatic challenges associated with this assay have hindered its widespread adoption in clinical diagnostics.

Here we present SpotWhisperer, a method that combines the accessibility of histopathology with the mechanistic insights of spatial transcriptomics and the convenience of chat-based data analysis. SpotWhisperer builds on recent advances in biomedical AI that infer spatial transcriptomics from H&E images and it interprets the resulting profiles using multimodal AI. Through this approach, SpotWhisperer provides the ability to analyze H&E images with natural-language queries, enabling the molecularly grounded interpretation of tissue composition and function at microscale spatial resolution.

We evaluated our method on a newly curated benchmark dataset for zero-shot H&E image property prediction with microscale resolution. The results demonstrate substantial performance gains over pathology VLMs, underscoring the benefit of integrating inferred spatial transcriptomic profiles into histopathology AI language models. Our benchmarking dataset and a video demonstration are provided on our project website http://spotwhisperer.bocklab.org.

## 2. Related work

The SpotWhisperer method builds on concepts introduced by VLMs for pathology, and extends their scope by incorporating microscale annotations with molecular information using multimodal models, which we introduce below.

### 2.1. Pathology vision-language models for H&E images

Pathology VLMs are trained on macroscale free-text annotations, which are sourced from clinical diagnostics and manually annotated by expert pathologists.

Most such models rely on multimodal contrastive learning (Radford et al., 2021) to connect images with their textual annotations, thereby enabling zero-shot classification across arbitrary free-text labels. Representative examples of this approach include PLIP (Huang et al., 2023) and CONCH (Lu et al., 2024), which we use here as baseline methods.

### 2.2. Spatial transcriptomics prediction

Spatial transcriptomics, i.e., the assessment of gene expression at genomic scale and spatial resolution, is commonly accompanied by H&E imaging of the same tissue slice. Consequently, a wealth of linked datasets at a microscale spatial resolution of 1 to 10 cells per spatial spot exists.

Based on these data, specialized machine learning models have been developed that can infer spatially resolved transcriptomic profiles from H&E images using convolutional neural networks (He et al., 2020), vision transformers (Pang et al., 2021; Jia et al., 2024; Zeng et al., 2022), and contrastive learning techniques (Xie et al., 2023).

The recent *DeepSpot* method (Nonchev et al., 2025) introduces two key advances. First, it leverages pathology foundation models trained on millions of H&E slides (Chen et al., 2024; Saillard et al., 2024; Filiot et al., 2024) to extract informative and robust representations from input image tiles. Second, it integrates multi-level tissue information covering zoomed-in and neighboring tissue morphology, in addition to the region of interest. This enables expression prediction of 5,000 genes — more than six times the number predicted by previous models — at improved accuracy.

### 2.3. Multimodal modeling of transcriptomes and text

Several works have linked transcriptomics to textual annotations, enabling zero-shot prediction with free-text labels.

Similar to established VLMs, LangCell (Zhao et al., 2024) and *CellWhisperer* (Schaefer et al., 2024a;b) employ multimodal contrastive learning (Radford et al., 2021) to link transcriptomes to text descriptions, facilitating efficient zeroshot prediction of cell type and other cell properties. Such models synergize with graphical single-cell applications, enabling interactive chat-based analysis of single-cell RNAseq data (Schaefer et al., 2024a;b).

## 3. Methods

SpotWhisperer advances computational pathology by augmenting H&E images with inferred molecular profiles and natural-language analysis at microscale resolution.

### 3.1. Microscale analysis of H&E images with natural language and molecular information

SpotWhisperer integrates the DeepSpot and CellWhisperer methods to enable molecularly informed analysis of H&E images with natural language (Fig. 1a) in four key steps.

**Figure 1.**
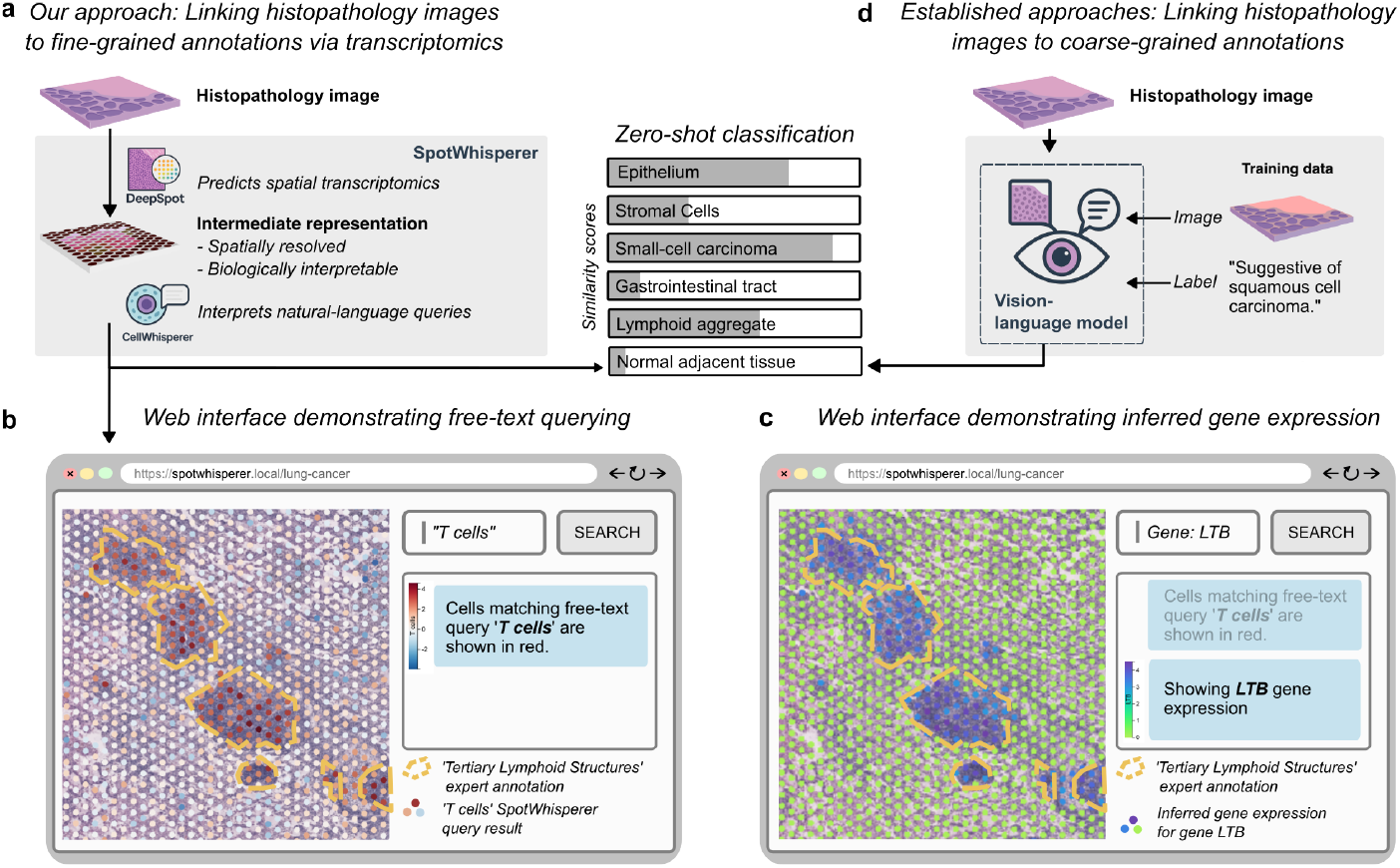
Zero-shot pathology image analysis at microscale and molecular resolution. (a) SpotWhisperer connects histopathological H&E images with fine-grained cell annotations via an interpretable gene expression embedding. (b) SpotWhisperer provides an interactive web UI for explorative H&E analysis using free-text natural language. (c) The SpotWhisperer web UI exploits inferred spatial transcriptomics. (d) Existing VLMs link H&E images to macroscale labels but ignore molecular information.

First, H&E images are subdivided into tiles centered on spot positions (see Appendix B). Second, DeepSpot (Nonchev et al., 2025) processes these tiles to predict spot-level (i.e., microscale) transcriptomic profiles. Third, the inferred spatial transcriptomics data are processed by the CellWhisperer embedding model (Schaefer et al., 2024b), yielding spot-level transcriptome embeddings. Fourth and finally, these embeddings are used for zero-shot inference with free-text labels, supported through CellWhisperer’s linked transcriptome and text embeddings.

To provide convenient access to our method, we extended the CellWhisperer web app (Schaefer et al., 2024b), facilitating intuitive annotation of H&E images using natural-language queries. This enables the direct search of cell types and states. For example, when searching for T cells in an H&E tumor image, our method clearly recapitulates expert annotations for “tertiary lymphoid structures” — regions that are known to harbor T cells and other immune cells (Fig. 1b). The molecular representation employed by SpotWhisperer facilitates biological interpretation of these predictions. Investigating the SpotWhisperer-identified spots, we identified strong expression enrichment for *LTB*, a gene known to be involved in the formation and maintenance of tertiary lymphoid structures (Fig. 1c). We provide a supplementary video of the analysis workflow on our project website.

SpotWhisperer facilitates molecular analysis and microscale annotation of H&E images, thereby supporting biological interpretation. It complements existing computational pathology models that focus on macroscale labels and do not link to the underlying molecular mechanisms (Fig. 1d).

### 3.2. Evaluation and benchmarking of spatially resolved natural language H&E image analysis

To quantitatively assess the performance of SpotWhisperer and future methods supporting microscale annotations, we compiled a benchmark dataset of H&E images with corresponding microscale ground-truth annotations, which we provide on our project website.

Our dataset is based on five published lung cancer tissue samples comprising a total of 16,032 patches imaged at 20× magnification (Dawo et al., 2025). We derived ground-truth annotations for each of these patches with two text-based labels: (i) a pathology-focused region annotation and (ii) a transcriptome-derived cell type annotation (Fig. 2).

**Figure 2.**
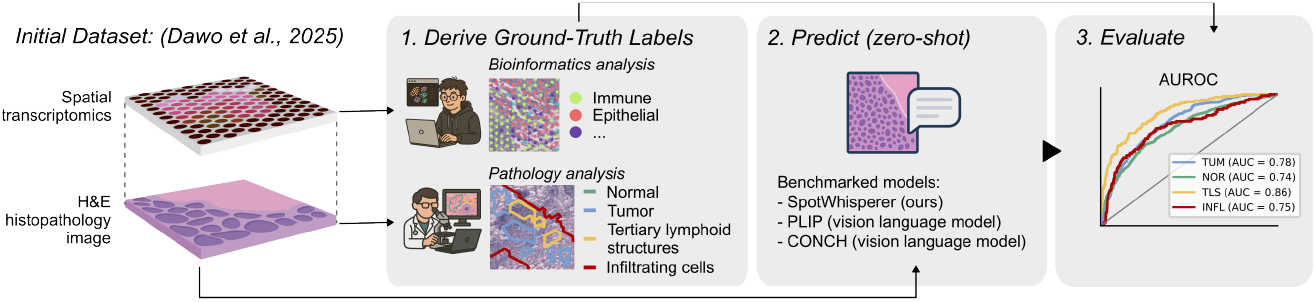
Schematic outline of evaluation dataset creation for the microscale H&E annotation benchmark and evaluation strategy of zero-shot H&E annotation models.

**Figure 3.**
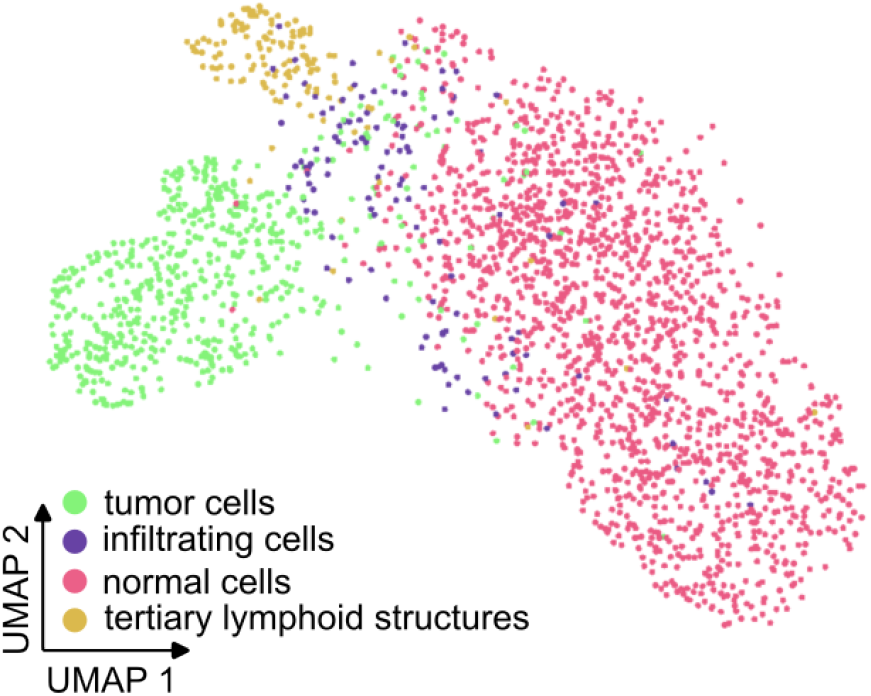
UMAP representing CellWhisperer embedding of 5000 HVGs inferred using DeepSpot on our lung cancer evaluation dataset with ground truth expert annotation labels.

We adapted the pathology-focused annotations from recent work (Dawo et al., 2025), where expert pathologists divided the H&E images into regions of *normal cells, tumor cells, infiltrating cells*, and *tertiary lymphoid structures*. These annotations were provided as abbreviations and translated by us into the natural language labels (see Table 3).

We derived the cell type annotations based on the spatial transcriptomics measurements provided alongside the H&E images in our evaluation dataset (Fig. 2) in two steps. First, we trained a logistic regression model on the comprehensive human lung reference atlas (Sikkema et al., 2023), considering only genes that were considered by the DeepSpot model. Then, we applied this model to the spatial transcriptomics data, transferring coarse-grained cell type labels *Immune, Epithelial, Endothelial, Stroma*, given that more specific cell type information was likely to be diffused by the multicellular resolution of the Visium technology.

Taken together, this dataset enables a data-driven evaluation of zero-shot H&E prediction methods with a focus on microscale and molecularly defined properties.

## 4. Results

To assess SpotWhisperer’s performance, we benchmarked it on our evaluation dataset in comparison to the state-of-theart VLMs PLIP and CONCH.

We performed zero-shot classification of spot-centered image patches (Fig. 1a) and compared them to the ground truth labels of our dataset (Fig. 2). We refer to Appendix B for details and to Fig. 1b for an example. Below we present the results for the two annotation types in the dataset.

SpotWhisperer was adapted to lung cancer tissue data by training a dedicated DeepSpot model (see Appendix A for details). We also investigated the potential to optimize the performance of VLMs through different patch sizes, but did not observe improvements (Appendix B.3).

### 4.1. Pathology-focused region prediction

On the pathology-focused region annotations, SpotWhisperer achieved the highest AUROC on three of the four region types and performed competitively on the fourth (see Table 1), highlighting the value of transcriptome-informed annotations for enhancing the accuracy of microscale histopathological analysis.

**Table 1.** AUROC scores and overall weighted AUROC of regionlevel annotation (mean (SEM)) for the evaluation dataset. Values in bold indicate the best-performing method.

| Region Type | PLIP | CONCH | SpotWhisperer (Ours) |
| --- | --- | --- | --- |
| infiltrating cells | 0.399 (0.036) | 0.550 (0.069) | **0.648 (0.089)** |
| normal cells | 0.453 (0.068) | 0.380 (0.057) | **0.658 (0.077)** |
| tertiary lymphoid structures | 0.881 (0.059) | **0.886 (0.031)** | 0.801 (0.050) |
| tumor cells | 0.627 (0.039) | 0.506 (0.048) | **0.760 (0.048)** |
| Overall | 0.554 (0.026) | 0.478 (0.025) | **0.717 (0.049)** |

### 4.2. Cell type prediction

Cell types are commonly defined by transcriptome signatures. They might thus be a challenging target for VLMs, but better detected by SpotWhisperer.

We benchmarked the three models in terms of their ability to predict corresponding labels in our evaluation dataset (see Table 4). SpotWhisperer achieved the highest mean AUROC, outperforming other models on three of the four cell types (see Table 2), underlining our method’s potential for predicting molecularly described annotations.

**Table 2.** AUROC scores and overall weighted AUROC of spotlevel cell type annotations (mean (SEM)) for the evaluation dataset. Values in bold indicate the best-performing method.

| Cell Type | PLIP | CONCH | SpotWhisperer (Ours) |
| --- | --- | --- | --- |
| Endothelial | 0.584 (0.025) | 0.527 (0.039) | **0.622 (0.048)** |
| Epithelial | 0.565 (0.025) | 0.549 (0.013) | **0.670 (0.052)** |
| Immune | 0.622 (0.022) | 0.564 (0.023) | **0.659 (0.035)** |
| Stroma | 0.521 (0.041) | **0.630 (0.069)** | 0.526 (0.060) |
| Overall | 0.586 (0.022) | 0.555 (0.009) | **0.656 (0.047)** |

**Table 3.** Queries used for region-level annotation.

| Expert Annotation | Query |
| --- | --- |
| TUM | tumor cells |
| NOR | normal cells |
| TLS | tertiary lymphoid structure |
| INFL | infiltrating cells |

**Table 4.**
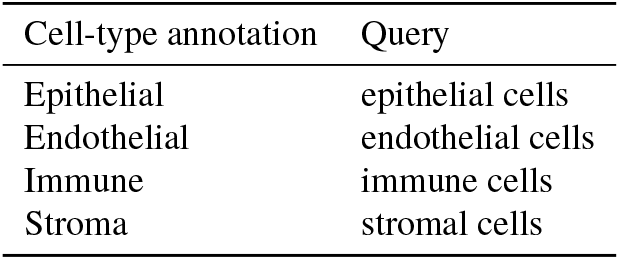
Queries used for spot-level annotations.

## 5. Discussion/Conclusion

With SpotWhisperer, we demonstrate the integration of spatially resolved molecular information into interactive histopathology analysis with free-text natural language queries. We employ transcriptome inference from H&E images to address the high cost and complexity of experimental spatial transcriptomics profiling, and we use a multimodal language model to enable chat-based interaction with such datasets, as illustrated in the demonstration video on our project website.

SpotWhisperer unlocks rich molecular annotations — as captured by CellWhisperer — for H&E image analysis using transcriptomic inference with DeepSpot. Our results demonstrate the downstream value for biological data interpretation, as SpotWhisperer outperforms state-of-the-art VLMs on pathology-focused and cell type annotation tasks. Nevertheless, VLMs were superior on a subset of tasks, including the identification of tertiary lymphoid structures (Table 1), indicating that integrating the macroscale strengths of VLMs with SpotWhisperer’s microscale focus could complement each other.

Our results and benchmark motivate further research on multimodal approaches for H&E analysis, including:

- Learning of tri-modal embedding space that jointly represents images, transcriptomics, and natural language.
- Increased spatial resolution of our benchmark and baseline method to the single-cell or subcellular level.
- Creation of a pan-tissue model, circumventing the need to train tissue-specific models.

Datasets, code, and access to our web UI will be provided upon archival publication of this work.

### 5.1. Study limitations

We point out several limitations of our study: First, we explored our approach on a single tissue type only, which we selected due to a sufficient availability of training data and the ability to curate independent evaluation data. We expect that newly published spatial transcriptomics data will soon facilitate the analysis across diverse tissue types. Second, we limited our evaluations to coarse-grained cell types such as stromal cells and epithelial cells due to the multicellular resolution of available transcriptomics data. Higher-resolution spatial transcriptomics data are rapidly becoming an established technology and will soon facilitate the assessment of fine-grained cell type predictions with our approach. Finally, while we ensured that baseline VLMs performed favorably on our chosen queries over rephrased alternatives, more elaborate methods such as prompt-ensembling could be explored for VLMs as well as for SpotWhisperer.

## A. SpotWhisperer implementation for lung cancer data

Spatial transcriptomics datasets have been generated and published for only a subset of human tissues. Here we trained DeepSpot in a tissue-specific manner focusing on lung cancer data matching the tissue type of our evaluation dataset. We further assessed the compatibility of DeepSpot-predicted transcriptomes with CellWhisperer (which is trained once and applied anywhere).

### A.1. DeepSpot model training

DeepSpot is trained as a regression model on the top 5000 highly-variable-genes (HVGs) in spatial transcriptomics data, using H&E image data as input features. For the lung cancer DeepSpot model, we used 10x Genomics Visium data from 36 lung cancer samples (De Zuani et al., 2024), which were generated independently from our evaluation data. From these data, the H&E slides are segmented into tiles overlapping with the spatial transcriptomics spot positions. For each spot, a bag of sub-tiles captures local morphology and a bag of neighboring spots incorporates the adjacent tissue environment. These image tiles are then processed to embeddings using H-optimus-0, a pre-trained pathology foundation model (Saillard et al., 2024), and serve as input to the trainable DeepSpot regression module. For a more detailed description of DeepSpot training, we refer to (Nonchev et al., 2025).

### A.2. Compatibility of DeepSpot predictions with CellWhisperer

To assess whether CellWhisperer meaningfully interprets DeepSpot-predicted spatial transcriptomics data, we embedded the inferred 5000 HVGs with CellWhisperer’s transcriptome embedding model and visualized the embeddings in a UMAP. We observed meaningful clustering by expert-level annotations, indicating that CellWhisperer recapitulated biological signals from the DeepSpot-predicted transcriptomics.

## B. Evaluating SpotWhisperer and vision-language models on lung cancer benchmark dataset

We evaluated SpotWhisperer, PLIP, and CONCH across five independent lung cancer samples (see Section 3.2) on spot-level annotations. Annotations comprise expert-annotated tissue labels (tumor, normal, tertiary lymphoid structure and infiltrating (immune) cells; see Table 3) and four bioinformatics-derived cell type labels (Epithelial, Endothelial, Immune and Stroma; see Table 4). For each of these two annotation groups, we calculated class probabilities using SpotWhisperer and the two VLMs, which we then derived AUROC scores from. This analysis was performed for each spot independently, where the spot positions were predefined by the source spatial transcriptomics data and the derived ground-truth labels. In use cases without such evaluation data, spot positions can be chosen freely, e.g. in a grid formation.

### B.1. Calculation of class probabilities with SpotWhisperer

To generate spot-level class probabilities, we embedded the class labels using CellWhisperer’s text embedding model and computed its cosine similarity with spot-level transcriptome embeddings, derived using CellWhisperer’s transcriptome embedding model, followed by softmax (Schaefer et al., 2024b).

### B.2. Calculation of scores with vision-language models

For the vision language models, PLIP and CONCH, we computed the similarity between the text queries and patches of the image localized at the annotated spots, as described in (Huang et al., 2023; Lu et al., 2024). Again, similarity scores were converted to probabilities using softmax. We note that CONCH was originally evaluated using prompt-ensembling to achieve its best performance (Lu et al. (2024), Extended Data Fig. 2). We opted not to report prompt-ensembling in our benchmark to better reflect a zero-shot setting where users provide a single query. This aligns with the broader aim of deploying models that are robust to variation in user queries without relying on prompt tuning or selection.

### B.3. Effect of patch size on vision-language model performance

Both PLIP and CONCH were trained on histopathology images that were often larger than the patch sizes corresponding to our microscale labels. We therefore tested whether larger patch sizes could improve the predictive performance of these baseline methods. To do so, we performed the same annotation task using a larger context area and taking the most frequent class in a given image patch as the overall label. We did not find a marked improvement in quality (Table 5).

**Table 5.**
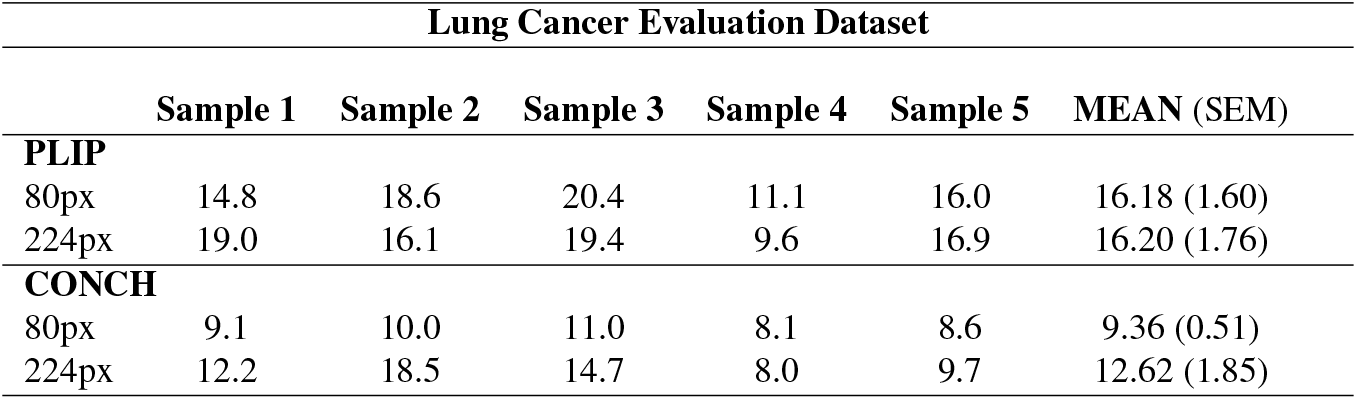
Difference in accuracy (%) on the region level-annotation task when VLMs are provided with a wider ‘context window’.

## Notes

### Competing Interest Statement

V.H.K. reports being an invited speaker for Sharing Progress in Cancer Care (SPCC) and Indica Labs; advisory board of Takeda; and sponsored research agreements with Roche and IAG, all unrelated to the current study. V.H.K. is a participant in a patent application on the assessment of cancer immunotherapy biomarkers by digital pathology; a patent application on multimodal deep learning for the prediction of recurrence risk in cancer patients, and a patent application on predicting the efficacy of cancer treatment using deep learning. G.R. is a participant in a patent application on matching cells from different measurement modalities which is not directly related to the current work. Moreover, G.R. is a cofounder of Computomics GmbH, Germany, and one of its shareholders. C.B. is a cofounder and scientific advisor of Myllia Biotechnology and Neurolentech which is not directly related to the current work.
The remaining authors declare no competing interests.

## References

Chen, R. J., Ding, T., Lu, M. Y., Williamson, D. F., Jaume, G., Chen, B., Zhang, A., Shao, D., Song, A. H., Shaban, M., et al. Towards a general-purpose foundation model for computational pathology. Nature Medicine, 2024.

Dawo, S., Nonchev, K., and Silina, K. 10x visium spatial transcriptomics dataset: Kidney (3) and lung (5) cancer with tertiary lymphoid structures. January 2025. doi: 10.5281/zenodo.14620362. URL https://doi.org/10.5281/zenodo.14620362.

De Zuani, M., Xue, H., Park, J. S., Dentro, S. C., Seferbekova, Z., Tessier, J., Curras-Alonso, S., Hadjipanayis, A., Athanasiadis, E. I., Gerstung, M., et al. Single-cell and spatial transcriptomics analysis of non-small cell lung cancer. Nature communications, 15(1):4388, 2024.

Filiot, A., Jacob, P., Mac Kain, A., and Saillard, C. Phikon-v2, a large and public feature extractor for biomarker prediction, 2024. URL https://arxiv.org/abs/2409.09173.

He, B., Bergenstråhle, L., Stenbeck, L., Abid, A., Andersson, A., Borg, å., Maaskola, J., Lundeberg, J., and Zou, J. Integrating spatial gene expression and breast tumour morphology via deep learning. Nature biomedical engineering, 4(8):827–834, 2020.

Huang, Z., Bianchi, F., Yuksekgonul, M., Montine, T. J., and Zou, J. A visual–language foundation model for pathology image analysis using medical twitter. Nature medicine, 29(9):2307–2316, 2023.

Jia, Y., Liu, J., Chen, L., Zhao, T., and Wang, Y. Thitogene: a deep learning method for predicting spatial transcriptomics from histological images. Briefings in Bioinformatics, 25(1):bbad464, 2024.

Lu, M. Y., Chen, B., Williamson, D. F., Chen, R. J., Liang,, Ding, T., Jaume, G., Odintsov, I., Le, L. P., Gerber, G., et al. A visual-language foundation model for computational pathology. Nature Medicine, 30(3):863–874, 2024.

Nonchev, K., Dawo, S., Silina, K., Moch, H., Andani, S., Consortium, T. P., Koelzer, V. H., and Raetsch, G. Deepspot: Leveraging spatial context for enhanced spatial transcriptomics prediction from h&e images. medRxiv, pp. 2025–02, 2025.

Pang, M., Su, K., and Li, M. Leveraging information in spatial transcriptomics to predict super-resolution gene expression from histology images in tumors. BioRxiv, pp. 2021–11, 2021.

Radford, A., Kim, J. W., Hallacy, C., Ramesh, A., Goh, G., Agarwal, S., Sastry, G., Askell, A., Mishkin, P., Clark, J., et al. Learning transferable visual models from natural language supervision. In International conference on machine learning, pp. 8748–8763. PmLR, 2021.

Saillard, C., Jenatton, R., Llinares-López, F., Mariet, Z., Cahané, D., Durand, E., and Vert, J.-P. H-optimus-0. 2024. URL https://github.com/bioptimus/releases/tree/main/models/h-optimus/v0.

Schaefer, M., Peneder, P., Malzl, D., Hakobyan, A., Sharma, V. S., Krausgruber, T., Menche, J., Tomazou, E., and Bock, C. Joint embedding of transcriptomes and text enables interactive single-cell RNA-seq data exploration via natural language. In ICLR 2024 Workshop on Machine Learning for Genomics Explorations, March 2024a. URL https://openreview.net/forum?id=yWiZaE4k3K.

Schaefer, M., Peneder, P., Malzl, D., Peycheva, M., Burton, J., Hakobyan, A., Sharma, V., Krausgruber, T., Menche, J., Tomazou, E. M., et al. Multimodal learning of transcriptomes and text enables interactive single-cell rna-seq data exploration with natural-language chats. bioRxiv, pp. 2024–10, 2024b.

Sikkema, L., Ramírez-Suástegui, C., Strobl, D. C., Gillett, T. E., Zappia, L., Madissoon, E., Markov, N. S., Zaragosi, L.-E., Ji, Y., Ansari, M., et al. An integrated cell atlas of the lung in health and disease. Nature medicine, 29(6):1563–1577, 2023.

Xie, R., Pang, K., Chung, S., Perciani, C., MacParland, S., Wang, B., and Bader, G. Spatially resolved gene expression prediction from histology images via bi-modal contrastive learning. Advances in Neural Information Processing Systems, 36:70626–70637, 2023.

Zeng, Y., Wei, Z., Yu, W., Yin, R., Yuan, Y., Li, B., Tang, Z., Lu, Y., and Yang, Y. Spatial transcriptomics prediction from histology jointly through transformer and graph neural networks. Briefings in Bioinformatics, 23(5):bbac297, 2022.

Zhao, S., Zhang, J., Wu, Y., Luo, Y., and Nie, Z. LangCell: Language-cell pre-training for cell identity understanding. arXiv [q-bio.GN], May 2024. URL http://arxiv.org/abs/2405.06708.

